# Uip4p modulates nuclear pore complex function in *Saccharomyces cerevisiae*

**DOI:** 10.1101/2021.09.02.458810

**Authors:** Pallavi Deolal, Imlitoshi Jamir, Krishnaveni Mishra

**Affiliations:** Department of Biochemistry, School of Life Sciences, University of Hyderabad, Hyderabad, India 500046

**Keywords:** Nuclear envelope, nuclear pore complex, yeast, Uip4

## Abstract

A double membrane bilayer perforated by nuclear pore complexes (NPCs) governs the shape of the nucleus, the prominent distinguishing organelle of a eukaryotic cell. Despite the absence of lamins in yeasts, the nuclear morphology is stably maintained and shape changes occur in a regulated fashion. In a quest to identify factors that contribute to regulation of nuclear shape and function in *Saccharomyces cerevisiae*, we used a fluorescence imaging based approach. Here we report the identification of a novel protein, Uip4p, that is required for regulation of nuclear morphology. Loss of Uip4 compromises NPC function and loss of nuclear envelope (NE) integrity. Our localisation studies show that Uip4 localizes to the NE and endoplasmic reticulum (ER) network. Furthermore, we demonstrate that the localization and expression of Uip4 is regulated during growth, which is crucial for NPC distribution.

## Introduction

Eukaryotes have intracellular compartmentalization and the biological processes are regulated spatially and temporally within membrane bound organelles. Nucleus is one of the most prominent organelles and an important defining feature of any eukaryotic cell. It harbours the genetic material within a double membrane bilayer that compartmentalizes it from the rest of the cellular components. The inner nuclear membrane (INM), acts as a site of regulation for various nuclear processes including transcription, mRNA export, ribosome biogenesis and DNA repair. Several INM associated proteins also provide structural support to the nucleus (Katta et al., 2014; Mekhail and Moazed, 2010). The outer membrane (ONM) is continuous with the endoplasmic reticulum which extends up to the cell periphery in yeast. The selective exchange of macromolecules between the nucleus and cytoplasm occurs through the macro-molecular assemblies called the nuclear pore complexes (NPCs). NPCs are embedded in the NE where the inner and outer nuclear membrane fuse. The estimated mass of an NPC in yeast is ∼60MDa comprising around 30 different types of nucleoporins (Beck and Hurt, 2016) with most of these present as 16 copies per NPC (Rout et al., 2000; Rajoo et al., 2018).

The NPC scaffold consists of three stacked rings, an inner ring that is sandwiched by two outer rings (cytoplasmic and nucleoplasmic), each of which has an eight-fold rotational symmetry (Joong Kim et al., 2018). The Nups are assigned to various classes primarily depending on their relative position in the entire complex (Joong Kim et al., 2018; Alber et al., 2007; Rout et al., 2000). The scaffold Nups (Yeast Pom152p, Pom34p and Ndc1) are the ones that nucleate the NPC assembly and hold the entire complex in place resulting in a cylindrical core (Onischenko et al., 2009; Yewdell et al., 2011). The phenylalanine-glycine (FG) repeat containing Nups are present along the inner side of the channel and are key regulators of the direction and flux of transport. The nucleoplasmic and cytoplasmic rings are concentric rings made of the 107-160 sub-complex (yeast Nup84 sub-complex). This comprises of Nup133p, Nup120p, Nup145Cp, Nup85p, Nup84p, Seh1p and Sec13p. Several nuclear and non-nuclear components are important in regulating the assembly, function and turnover of NPCs (Allegretti et al., 2020). The ONM is known to share several proteins with the endoplasmic reticulum (ER). Some of these ER proteins, such as the reticulons-Rtn1 and Yop1, and membrane metabolism related proteins-Brr6, Brl1 and Apq12, also contribute to NPC biogenesis and assembly (Casey et al., 2015; Dawson et al., 2009; Scarcelli et al., 2007; Lone et al., 2015).

NPC assembly is known to operate via two major pathways (Fernandez-Martinez and Rout, 2009; Otsuka et al., 2018). One is *de novo* assembly during interphase when nucleoporins are recruited to a pore site at nuclear membrane. Second, post-mitosis, when sub-complexes and membrane vesicles are recruited to NE during nuclear envelope reformation. In organisms that undergo closed mitosis, only the former mechanism takes place. Defective assembly of NPC intermediates and dysregulation of nuclear quality control pathway results in abnormal nuclear shape (Mészáros et al., 2015; Webster et al., 2014; Deolal and Mishra, 2021). Mis-localization and altered stoichiometry of nucleoporins has been also linked to aging and clinical pathologies (Sakuma and D’Angelo, 2017). NE extension or herniations with NPC clustering have been observed in yeast as a result of failed nuclear protein quality control pathways (Wente and Blobel 1993, Webster et al 2016, Thaller et al 2019). In most of the cases where NPC aggregation is seen, accompanying nuclear shape changes are also observed (Wente and Blobel 1993).

Presence of nups outside of the NE is atypical but cytosolic spots containing NPC components have been reported under various physiological and non-physiological conditions in budding yeast (Colombi et al., 2013; Makio et al., 2013; Webster et al., 2014; Lord and Wente, 2020). While some of the NPC components might assemble in the cytosol, they are not recruited at the NE until the transmembrane and scaffold nups have created pore site for insertion. In yeast some of these cytoplasmic spots are pools of nups and remain associated with lipid droplets (Kumanski et al., 2021). While in other cases, these components are retained to prevent incorporation of damaged nups at the NE thereby protecting NE integrity (Webster et al., 2014). Cytosolic spots of nups have been observed in yeast mutants that affect the assembly and stability of NPCs (Flemming et al., 2009; Onischenko et al., 2009).

In order to identify novel components involved in maintaining nuclear architecture we initiated a fluorescence microscopy based genome-wide screen approach in budding yeast. Yeast serves as a good model for such a screen because unlike metazoans it undergoes closed mitosis and the yeast nuclear membrane does not disassemble during mitosis (Rout and Wente, 1994). We found that the loss of certain physical interactors of Ulp1, an inner nuclear membrane associated deSUMOylase, resulted in nuclear shape change and also affected distribution of nups. Loss of one of the proteins, namely Uip4, compromised the nuclear permeability barrier. We show that Uip4 localizes predominantly to the NE/ER and loss of Uip4 results in dramatic mis-localization of the NPCs. Furthermore, we also find that Uip4 expression is increased when the cells transition to stationary phase of growth. Despite the knowledge of basic structural components of NE and NPCs, a clear understanding of the underlying mechanisms that contribute towards the maintenance of shape of NE and integrity of associated complexes is lacking. This work opens up avenues for understanding multiple ways in which cellular components can contribute towards maintenance of the nuclear shape, integrity and thereby function.

## Materials and Methods

### Yeast strains and growth

The yeast strains used in this study are listed in table 1. All strains were grown either in YPD (1% yeast extract, 2% peptone and 2% dextrose) or SC (0.75g SC dropout mix, 2% dextrose) media. Yeast transformation was performed using standard lithium acetate based protocol. For all experiments, yeast strains were grown to mid-log phase at 30ºC. For stationary phase, cells were harvested after 36 hours of growth. C-terminal 13MYC tag at the genomic loci of *UIP4* and *ESC1* was introduced by PCR based homologous recombination using pFA6a-13MYC-HIS3MX6 as described in (Longtine et al., 1998). The high-efficiency transformation of PCR product for deletion and tagging was performed as described (Gardner and Jaspersen, 2014).

**Table 1:**
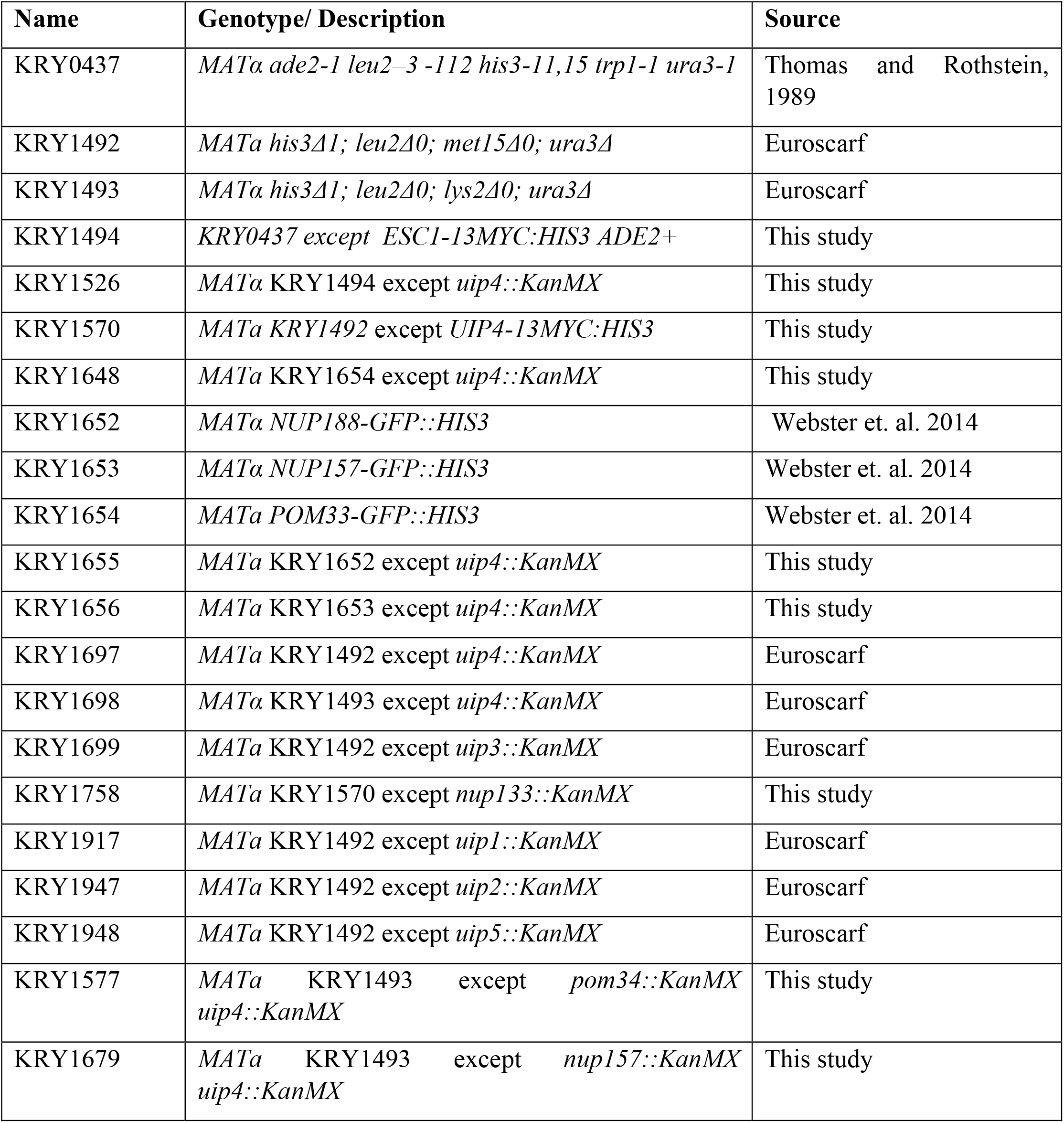
List of strains used in the study.

### Plasmids

All plasmids used in this study have been described in table 2. GFP-Esc1 (Male et al., 2020), GFP-Nup49 (Bucci and Wente, 1998) and NLS-2XGFP (Stade et al., 1997) have been previously described. To generate pRS313-UIP4, the gene was amplified from genomic DNA of wild type BY4741 strain with its own promoter (500bp upstream of start) and 3’UTR (150 bp downstream of STOP) using Vent DNA Polymerase. For over expression of UIP4, the gene was amplified beginning from start codon upto 150 bp downstream of STOP. The fragment was inserted into the BamHI/SalI site of pBEVY-L under the *pGPD* promoter. All constructs were confirmed by sequencing.

**Table 2:**
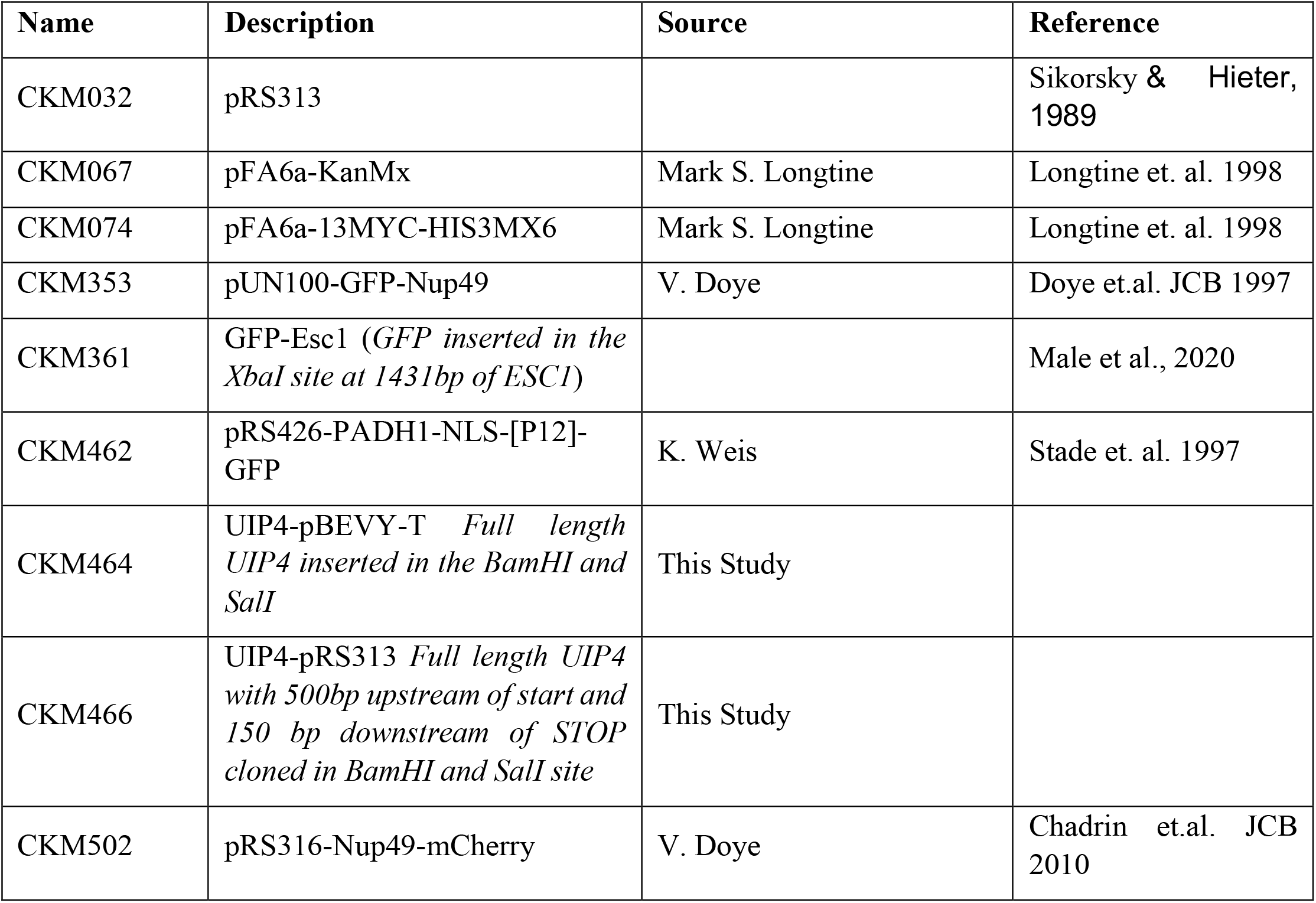
List of plasmids used in the study.

### Live cell imaging

Cells grown were harvested by centrifugation and the pellet was washed twice with fresh SC media. For images shown in Fig1A, 1B, S1C and S2I, cells suspended in SC media were added to Concanavalin A coated slides. The cells were allowed to settle for 5 minutes and excess liquid was removed using a vacuum pump. For nuclear staining, 5ul of 50ng/ml DAPI was added to the cells in dark. After 5 minutes, 4ul of Slow Fade Anti-fade mounting medium (Invitrogen S36936) was added and coverslip placed. The edges were sealed gently using nail polish. For other live cell and time lapse imaging, cells resuspended in growth media were mounted onto a 35mm cover glass bottom dish coated with ConA. After allowing the cells to settle, unbound cells were rinsed with SC media. Images were acquired in Leica TCS SP8 equipped with a temperature controlled stage set at 30ºC. Cycloheximide was added to a final concentration of 10µg/ml in the imaging medium for experiment shown in Fig S2D.

**Figure 1:**
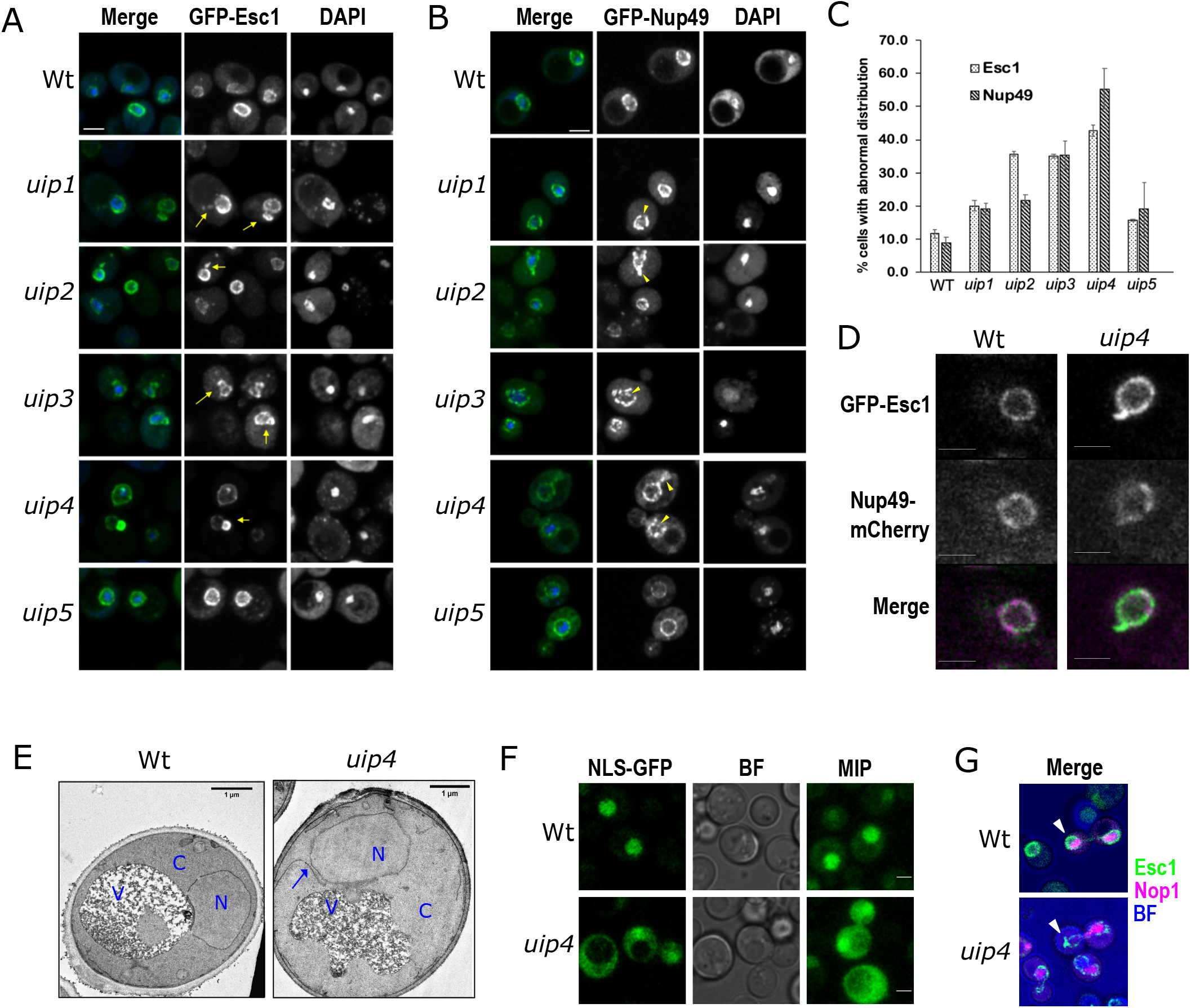
Loss of UIPs results in altered nuclear shape and nuclear import. **A, B**. Wt and strains harboring indicated deletions were examined for nuclear morphology and distribution of nuclear pores using GFP-Esc1 (A) and GFP-Nup49 (B) plasmids respectively. Maximum intensity projection (MIP) of representative cells are shown. DAPI staining is used to define the nucleus. Scale-2µm **C.** Quantification of defects is shown in the bar graph as the fraction of cell population showing abnormality (n>300, 3 independent experiments, error bars indicate SEM). **D.** Mid-focal planes of Wt and uip4 co-transformed with GFP-Esc1 and Nup49-mCherry are shown. Scale-2µm. **E.** Transmission electron micrographs for Wt and uip4 are shown. N-nucleus, C-cytoplasm, V-vacuole. Arrow-NE extension in uip4. Scale-1µm **F.** Nuclear import was tested in Wt and uip4 bearing NLS-2X GFP plasmid. Scale-2µm **G.** MIP of dividing Wt and uip4 cells with NE (GFP-Esc1) in green and nucleolus (mRFP-Nop1) in magenta are shown. Arrow heads-associated NE abnormality.

### Immunofluorescence

Immunofluorescence was performed as described in (Tirupataiah et al., 2014). Briefly, cells were fixed in 3.7% formaldehyde at 30ºC for 20 minutes, followed by three washes with sterile distilled water. Cells were then treated with 10mM DTT and 0.1M EDTA-KOH for 10minutes at 30ºC. Spheroplasts were generated by treating the cells with 0.25mg/ml Zymolyase 100T at 30ºC until the cell wall was digested. Digestion of cell wall was confirmed by visualising a drop of cell suspension under a light microscope. Appropriate amount of spheroplasts were dropped onto a poly-L-Lys coated slides. The cells were permeabilized by treating with methanol and acetone for 5minutes and 30 seconds respectively. Blocking was done using 1% BSA prepared in 1X-PBS (pH 7.4) with 0.1% Tween-20. Primary antibodies α-Myc (Abcam-9E11) and αNsp1 (Abcam ab4641) were used at a dilution of 1:800, and α-Pdi1(Abcam ab4644) at 1:500 and stained overnight at 4ºC in a closed, moist slide box. Staining with secondary antibodies labelled with either AF488 (Invitrogen) or Cy3 (Jacksons) at a dilution of 1:1000 was done at room temperature for 2hrs. DAPI staining was done for 5 minutes followed by mounting with Prolong gold antifade mountant (Invitrogen, P36930).

### Electron Microscopy

For transmission electron microscopy, yeast cell cultures were grown in SC media and harvested at mid-log phase (0.8 OD at A600). For detailed method refer to Male et al., 2020. Briefly, cells were fixed in ice-cold fixing solution (2% glutaraldehyde and 1% paraformaldehyde, 1mM MgCl2, in 50 mM potassium phosphate buffer (pH 6.8) for 2 hrs at 4°C, followed by post-fixation in 4% KMnO4 for 1 hr at room temperature (RT). Cells were treated with freshly prepared 2% uranyl acetate for 1 hr at RT and subsequently washed. Dehydration of cells was followed by clearing with propylene oxide and infiltration with the combination of 100% propylene oxide and 100% spur low viscosity resin (TED PELLA.INC, USA).

### Western Blotting

Cells transformed with desired plasmid were grown in selection medium either upto mid-log phase or stationary phase, as desired. Proteins were extracted using TCA method of protein precipitation (Abraham et al., 2019) Briefly, 200µl of 20% TCA was added to the cell pellet and were lysed by adding glass beads and vortexing at high speed for 3 minutes. The lysate was transferred to a fresh 1.5ml tube. The tube containing glass beads was washed twice with 200µl of 5% TCA and vortexing for 1 minute. The lysate was collected and added to the previous tube. The entire lysate was spun at 13000rpm for 10 minutes. Supernatant was discarded completely and the pellet was dissolved in 200µl of Laemmli buffer. 2M Tris was added drop-wise until the buffer turned blue. Proteins were denatured by boiling in Laemmli buffer for 5 minutes at 95ºC. SDS-PAGE was done followed by a standard semi-dry method of transfer to PVDF membrane for western blotting. Primary anti-Myc (Abcam 9E11, 1:10000), anti-GFP (Abcam ab290, 1:3000), anti-Nsp1(Abcam ab4641, 1:10000) and anti-actin (Santa Cruz sc-47778, 1:5000) were detected by HRP tagged secondary anti-rabbit (Abcam ab97051) and anti-mouse (Santa Cruz sc-2005). The signal was detected by ECL reagent (GBiosciences) and imaged in ChemiDoc Imaging system by BioRad.

### Quantification

Quantification of the phenotype for all presented data was done from at least three independent experiments. At least 100 cells were analysed from each experiment. Representative images are shown. Bar graphs represent the fraction of population showing the indicated phenotype. Where shown, error bars represent SEM. p-values to assess significance was determined by Student’s t test. NPC aggregation index was determined as described in (Lord and Wente, 2020). Briefly, the NE outline based on the signal from indicated nup was traced in FIJI using the freehand tool. The selection was converted to a line and fluorescence intensity values were obtained using the plot profile function. The intensity values for ∼20-25 cells from at least two independent experiments were copied to excel sheet and respective background fluorescence intensity was subtracted. Next, for each NE, average intensity was subtracted from the individual values and the absolute difference was obtained. The standard deviation of this difference was divided by the average intensity of each NE and referred to as the aggregation index.

## Results

### Loss of Ulp1 interacting proteins results in distorted nuclear morphology

In order to identify the components that contribute to the process of regulating nuclear shape, we initiated a fluorescence microscopy-based screen (Male et al., 2020; Deolal and Mishra, 2021 and unpublished data). To this end, non-essential gene knock-out strains of BY4741 background were systematically transformed with a GFP-tagged inner nuclear membrane protein (ScEsc1). We have previously validated the use of GFP-Esc1 as a marker for nuclear shape in budding yeast (Male et al., 2020). Wild type cells display a round, nuclear envelope staining (Fig1A, row1), but several gene knockouts identified in the screen have abnormal nuclear shape (Male et al., 2020; Deolal and Mishra, 2021). While screening the non-essential gene knockout collection of *S. cerevisiae* for nuclear shape in a chromosome-wise manner, we found that loss of *UIP2* and *UIP3* from chromosome I resulted in altered nuclear shape (Fig.1A). As both *UIP2* and *UIP3* are interactors of Ulp1p, we drew out knockouts of other non-essential <u>U</u>lp1 interacting <u>p</u>roteins, Uip1 –Uip5, to screen for nuclear shape defects (FigS1A). UIPs are proteins that were identified in a two-hybrid screen for interactors of the deSUMOylase, Ulp1p (Takahashi et al., 2000a). Uip1p and Uip6p are established components of NPCs in yeast; Uip1p is a non-essential FG nucleoporin-Nup42 and Uip6p is an essential cytoplasmic Nup-Gle1 (Rout et al., 2000). The shape of the nucleus as marked by GFP-Esc1 was severely distorted upon loss of *UIP3* and *UIP4* (Fig1A-arrows, 1C). Clear membrane extensions and flares could be seen upon loss of *UIP*2. *uip3*Δ and *uip4*Δ cells had large membrane blebs. Previous studies have reported an intricate dependence of nuclear shape and NE dynamics on the distribution of NPCs (Schneiter and Cole, 2010). Mutants that have aberrant NPC distribution are often accompanied by nuclear shape abnormalities. Also, Esc1 is known to modulate the Ulp1 distribution and nuclear basket assembly at the NE (Lewis et al., 2007). Therefore, we wanted to see if the NPC distribution is affected upon loss of these *UIPs*. GFP-Nup49 was used to examine the distribution of NPCs. We found that loss of UIPs resulted in clustering of Nup49 as compared to its distribution in wild type cells (Fig1B-arrow heads, 1C). This clustering phenotype was very prominent in a large fraction of *uip4*Δ cells (Fig1C). A large fraction of *uip4*Δ cells had severe nuclear shape abnormalities as well. Blebs and extensions of nuclear membrane was the most prominent nuclear shape defect in *uip4*Δ (Fig1D). We further assessed the nuclear shape abnormality in *uip4*Δ by transmission electron microscopy. The micrographs representing the quasi-round nuclear shape in wild type and abnormal nuclear shape with extensions in *uip4*Δ are shown in Fig1E.

### Uip4 is required for maintaining nuclear structure and function

NE serves as a selective barrier between cytoplasm and nucleoplasm. In order to check the integrity of NE and functional competence of the nuclear pores, we tested the nuclear import capability of cells lacking UIPs using NLS-GFP construct (Stade et al., 1997). Wild type cells accumulate NLS-GFP in the nucleus efficiently as evidenced by a bright nuclear fluorescence (Fig1F, S1B). Similarly, *uip1Δ, uip2Δ, uip3*Δ and *uip5*Δ cells also import NLS-GFP in to the nucleus (FigS1B). However, loss of *UIP4* showed a large defect in the nuclear import of NLS-GFP (Fig1F, S1B). This indicates a compromised nuclear permeability barrier upon loss of Uip4 and suggests that the clustered pores in other UIPs are functionally competent for nuclear import unlike in *uip4*Δ.

Despite compromised nuclear structure, none of the deletion mutants had any growth defects (S1C). Among the 5 UIPs studied for nuclear defects, loss of *UIP4* resulted in the most severe phenotype of a compromised nuclear structure and function. Therefore, we wanted to further delineate the effect of Uip4 and characterize this protein. Defective nuclear shape has been associated with delayed mitosis (Webster et al., 2009). Therefore we sought to monitor the nuclear shape dynamics during cell division by live cell microscopy. Cells were grown in a 35mm cover glass bottom dish in the presence of SC medium at 30ºC. *S. cerevisiae* undergoes closed mitosis therefore in wild type cells the nuclei remained mostly round-elliptical with a uniform distribution of Esc1p around the nuclear periphery (Fig1G). On the other hand, *uip4*Δ display a wide range of nuclear morphologies during nuclear transmission to the bud (Fig 1G, FigS1D, arrow heads). This resulted in an increased nuclear division time for *uip4*Δ as compared to the wild type (FigS1E). However the overall growth rate for wild type and *uip4*Δ was not distinguishable (Fig S1F).

### Loss of Uip4 results in abnormal NPC distribution

Proper NPC assembly and distribution is crucial for maintenance of the nuclear shape (Wente and Blobel, 1993; Titus et al., 2010). While in wild type cells, less than 10% cells had nuclei that were not spherical, in *uip4*Δ, over 50% of cells had distorted nuclear shapes and clustered nuclear pores (Fig1C). We quantified the clustering of NPCs by calculating the aggregation index based on the distribution of fluorescence signal intensity of GFP tagged nucleoporins along the NE. The aggregation index for GFP-Nup49 in *uip4*Δ cells was found to be significantly higher than that in wild type cells (Fig2A, 2B). We then examined the distribution of inner ring nucleoporins namely Nup157 and Nup188. Both of these nups were also clustered upon loss of *UIP4* (Fig2A, 2B). The transmembrane nup, Pom33, showed a differential localisation in the *uip4*Δ cells. A minor fraction of Pom33 is localized to the ER in addition to the NE in wild type cells (Chadrin et al., 2010). Apart from clustering at the NE, there was a reduction in the distribution of Pom33 to the cortical ER in *uip4*Δ as compared to wild type cells, with most of Pom33 present in the NE (Fig2C). These observations were also reiterated by indirect immunofluorescence using GFP antibody to detect Nup157, Nup188 and Pom33, and co-stained with anti-Nsp1 (Fig S2A). Loss of Uip4 therefore seems to affect distribution of multiple nups.

**Figure 2:**
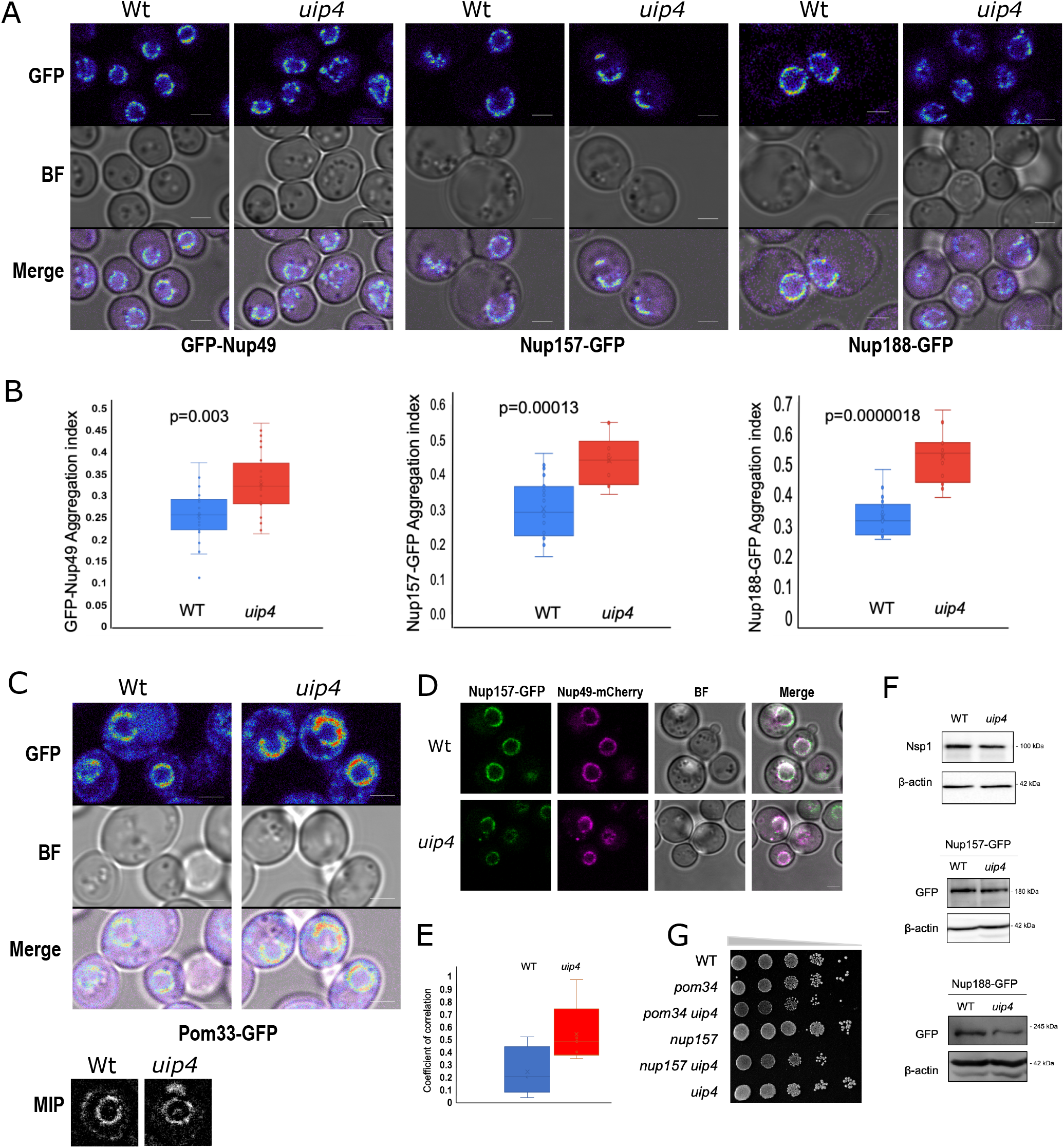
Loss of UIP4 results in NPC clustering. **A.** Live cell imaging was done for Wt and uip4 strains expressing indicated GFP tagged Nups from their endogenous loci. Spectrum lookup table is used for GFP signal (red-max, blue-min). Scale-2µm **B.** The box plot represents the aggregation index calculated from GFP signal along the NE in the mid-focal plane of 25-30 individual cells from Wt (blue) and uip4 (red). Horizontal line is the mean value. **C.** Live cell imaging was done for Wt and uip4 strains expressing Pom33-GFP. Spectrum lookup table is used for GFP signal (red-max, blue-min). Scale-2µm The inset shows MIP of a representative nucleus from Wt and uip4 in grayscale. **D.** Wt and uip4 co-expressing Nup157-GFP and Nup49-mCherry are shown. Scale-2µm. **E.** The extent of co-localization for Nup157-GFP and Nup49-mCherry shown in D is assessed by measuring the correlation coefficient between the two signals. Horizontal line represents the mean. **F.** Western blot analysis was done to compare the expression levels of Nsp1, Nup157 and Nup188 in Wt and uip4 cells harvested from mid-log phase. α-GFP was used to detect endogenously tagged Nup157 and Nup188 and Actin is the loading control. **G.** Overnight cultures of the indicated strains were taken and sub-cultured by inoculating equal number of cells in a fresh medium for 4 hours. Cells were then harvested and 10-fold dilutions were serially spotted on a SC-plate. The plates were incubated at 30ºC for 2 days prior to imaging.

Apart from aggregation of NPCs at the NE, we also observed presence of cytosolic spots of nups in *uip4*Δ (Fig2D). Although cytosolic pool of nucleoporins in wild type cells has been reported earlier (Colombi et al., 2013; Makio et al., 2013; Kumanski et al., 2021), this fraction of cell population was at least three-fold higher in *uip4*Δ cells (FigS2B). Complementation with a plasmid borne copy of *UIP4* expressed from endogenous promoter could rescue the phenotype in *uip4*Δ (FigS2B). We assessed the relative distribution of nucleoporins from two sub-complexes between wild type and cells lacking UIP4. The NPC clusters at the nuclear periphery do not differ in their relative nup constitution between wild type and *uip4*Δ as determined based on the degree of colocalization of Nup157 and Nup49 (Fig2D, S2C). Contrary to this, the cytosolic spots observed in *uip4*Δ contain more than one nup unlike in wild type (Fig2E, S2C).

The cytosolic foci observed in *uip4*Δ could be either assembled NPC assembly intermediates or disintegrated NPCs. In order to delineate if the cytosolic signal is indicative of *de novo* NPC assembly intermediates, we treated wild type and *uip4*Δ cells with cycloheximide, a protein translation inhibitor. We monitored GFP-Nup49 in wild type and *uip4*Δ cells by time-lapse live cell imaging after addition of cycloheximide (S1D). We did not see any increase in the cytosolic foci indicating that these cytosolic spots are intermediates of NPC assembly (Fig S2D). Interestingly, we also observed that after two hours of cycloheximide exposure, even in wild type cells, the NPCs began to cluster and nuclear morphology also began to distort (FigS2C, S2E). The effect of loss of *UIP4* on the stability and turnover of individual nucleoporins warrants further investigation. Our western blot results indicate that the total protein levels for Nsp1and Nup157 do not differ much between wild type and the *uip4*Δ and the modest reduction seen in Nup188 is also statistically insignificant (Fig 2F), again indicating that these are unlikely to be disintegrating NPCs.

Since loss of *UIP4* compromised function of NPCs, we wanted to test if Uip4 interacts with nucleoporins or other NE proteins that contribute to NPC stability. To see if there is a genetic interaction, we created double mutants with POM34 and NUP157. Pom34 is one of the three primary transmembrane nup in yeast (Strunov et al., 2011). Pom34 is important during the NPC biogenesis whereas Nup157 is required for ensuring proper assembly of pores at later stages (Makio et al., 2009). We did not observe a strong genetic interaction of *UIP4* with these nups (Fig2G). Further, we attempted to test if the effect of Uip4 on NPC function is due to its interaction with Ulp1 (Takahashi et al., 2000b; Rouvière et al., 2018). Although Uip4 was originally identified in a yeast two-hybrid screen for interactors of Ulp1, we did not find any positive interaction between Ulp1 and Uip4 by direct two-hybrid (data not shown). Additionally, in a yeast two-hybrid screen to obtain other physical interactors of Uip4, we did not detect any nucleoporin or nuclear protein that is known to directly affect NPC function. Taken together, these results indicate that although the presence of Uip4 contributes positively towards NPC assembly and stability, this effect is likely to be a consequence of either a transient association of Uip4 with nups or an indirect association.

### Uip4 localises to NE/ER

Uip4 harbors an N terminal Early Set glycogen binding domain (1-87aa) and a MDN1 superfamily midasin domain (73-271aa) (Marchler-Bauer et al., 2011) (Fig S2F). It lacks a transmembrane membrane domain but has a KKXX (_277_KKLL_280_) motif, suggesting potential ER membrane retention. In order to study the expression and subcellular localization of Uip4p, *UIP4* was tagged with 13xMyc epitope (Longtine et al., 1998). Tagging Uip4 did not cause a loss of function as the nuclear shape and NPC distribution in strain expressing 13xMyc tagged UIP4 was similar to that of wild type cells (FigS2G). To test the localization of Uip4, we performed indirect immunofluorescence using anti-myc antibody. We co-stained the cells with Pdi1 as an ER marker. In consonance with the presence of KKLL motif, indirect immunofluorescence results showed that Uip4 localizes to two prominent rings resembling ER distribution, in addition to some cytosolic signal (Fig3A, FigS2H). The distribution of Uip4-13xMyc largely overlapped with Nsp1 at the perinuclear ER suggesting that Uip4p is an NE/ER protein (FigS2H). Expression of the protein was confirmed by western blot analysis (Fig3B). We observed a higher expression of Uip4 in cells harvested from stationary phase as compared to those in mid-log phase. The Uip4p localization was largely unaffected during stationary phase (Fig3C), although, a more continuous staining of Uip4 was present at the NE/ER. Wild type cultures from stationary phase of growth also have cytosolic presence of nups that display foci like distribution distinct from the NE (Fig3C, arrow). The Uip4 signal did not overlap with the cytosolic spot of Nsp1. To test whether Uip4p localization is affected by strong clustering of NPCs, we tested the localization of Uip4p in *nup133*Δ, a strain which is known to have clustered NPCs (Doye et al., 1994). We found that in *nup133*Δ, Uip4 was localized extensively to ER (Fig3D). Even in this case the two signals do not co-localize, suggesting that pore clustering does not influence Uip4p localization and Uip4p does not physically associate with NPCs.

**Figure 3.**
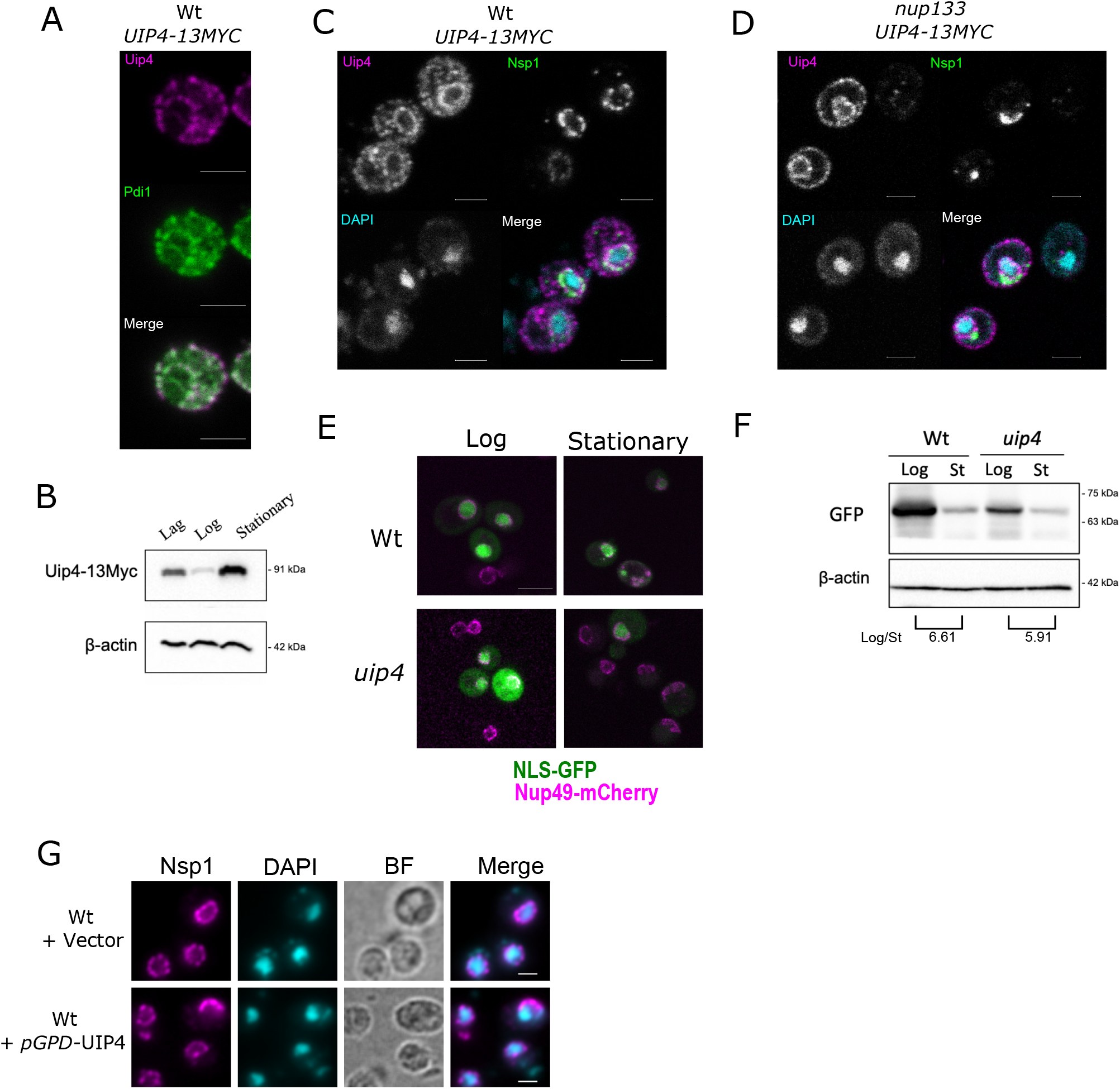
Uip4 localises to NE/ER and its overexpression exacerbated NE defects. **A.** Indirect immunofluorescence using α-myc was performed in the strain encoding UIP4-13xMyc to check the localization of Uip4. Co-staining with an ER marker Pdi1 is shown. Scale-2µm **B.** The western blot is representative of the expression level of Uip4 in cells harvested from either lag (O.D.600-0.2-0.4), mid-log (O.D.600-0.8-1.2) or stationary phase (O.D.600-3.0-3.5). Uip4 was detected using α-myc. Actin is used as a loading control. **C.** Co-staining of Uip4 with a NPC marker, Nsp1, was done in cells harvested from stationary phase. Scale-2µm **D.** Localization of Uip4 in nup133Δ background was checked. Co-staining of Uip4 with a NPC marker, Nsp1, is shown. Scale-2µm **E.** Live cell imaging was performed in Wt and uip4 cells bearing NLS-GFP plasmid harvested from either mid-log or stationary phase. The overlay images showing NLS-GFP (green) and Nup49-mCherry (magenta) are indicative of the observation. Scale-5µm **F.** Western blot showing protein level of NLS-GFP detected using α-GFP is shown for wild type and uip4 cells harvested from log and stationary (St) phase. The quantification of the NLS-GFP in Log/Stationary phase is shown (Average of 2 independent experiments). Actin is used as a loading control. **G.** The distribution of NPCs upon UIP4 OE was checked by monitoring localization of Nsp1, an FG nucleoporin. Indirect immunofluorescence was performed in WT cells transformed with either E-pBevyL or pGPD-UIP4-pBevyL. Arrow head indicates non-nuclear spots of Nsp1. Scale-2µm

### Regulated Uip4 expression is critical for NPC function

We found the Uip4p levels to be lowest during log phase when the cells are actively dividing and highest during the stationary phase when the growth is ceased (Fig3B). This expression correlated with the higher defect in NPC distribution in *uip4*Δ cells during stationary phase (FigS2I). Similar to the cells from log phase culture, we observed more cells showing cytosolic foci of Nup49 in stationary phase as well, for *uip4*Δ. This indicates that Uip4 could be important for NPC distribution particularly during stationary phase. To assess this further, we looked at the NE integrity by import of NLS-GFP (Fig3E). Fluorescence imaging indicated an overall reduction in the signal intensity for cells harvested during stationary phase for both wild type and *uip4*Δ (Fig3E). Western blots also confirmed the lower level of NLS-GFP expression in both cells harvested from stationary phase as compared to those harvested in log phase (Fig3F). In addition to defective nuclear import, in *uip4*Δ cells the expression level of NLS-GFP was lower than respective wild type in all stages of growth (Fig1F, Fig3E, Fig3F). Importantly, in the stationary phase, the reduced NLS-GFP signal was almost entirely nuclear in wild type cells. But in *uip4*Δ the NLS-GFP signal was either from dead cells or whole cells, and hardly any cells showed nuclear accumulation, indicating a more compromised NPC function.

Having established the physiological conditions that regulate Uip4 expression, we tested the NPC distribution upon ectopic overexpression of Uip4. We overexpressed Uip4 from a strong promoter and observed severe clustering of Nsp1 at the NE (Fig3G). Cytoplasmic spots were also observed upon Uip4 overexpression (Fig3G). Nuclear accumulation of NLS-GFP was also greatly reduced suggesting that the nuclear import is severely defective when Uip4 is overexpressed (FigS2J). We found that the NPC localization and NE integrity defects were exacerbated when Uip4 was overproduced. As both loss and overproduction of Uip4 led to distorted nuclear envelope architecture and defective protein import function, it appears that the levels of Uip4p are important for regulating its function at the nuclear envelope. Taken together, these results suggest that the dosage and distribution of Uip4 is critical for its function in budding yeast.

### Summary

We have identified a novel role of a previously uncharacterised protein-Uip4 which localises to NE/ER. While a comparative sequence analysis approach has shown that Uip4 is an ascomycete specific protein (Garapati and Mishra, 2018); proteins with E-Set domain are known in all eukaryotes. Therefore organisms could have functional homologs of Uip4 that could not be identified by sequence comparisons. We find that loss of Uip4 leads to assembly of functionally compromised NPCs. It is likely that altered Uip4p regulation affects the quality of NE by causing dysregulation of NPC stoichiometry. Uip4 could be involved in stabilizing the assembly intermediates of NPC or the entire complex itself at the NE by virtue of its membrane association. The amphipathic helix at the C-terminal end of Uip4 is likely to act as its in-plane membrane anchor (predicted by AMPHIPASEEK-Combet et al., 2000). In the absence of *UIP4*, we speculate that the incorporation of these substrates into the NE/NPC is attenuated and therefore can be seen in cytosol.

Even though Uip4 was not found to be present physically at the clustered pores, our results show that presence of Uip4 is required for proper distribution of the pores. This dependence could be due to either a temporary association or an indirect interaction. Many proteins with other primary functions are known to affect NPCs due to their transient association with nups or dynamic localization to nucleus (Aitchison and Rout, 2012; Scarcelli et al., 2007; Casey et al., 2015). Since we find that Uip4 is overexpressed during stationary phase in wild type cells, it would be interesting to study the molecular mechanism of Uip4 induced NPC aggregation. Compared to log phase cells, a larger fraction of Uip4 is recruited to NE in stationary phase and also in *nup133*Δ (compare Fig3A to Fig3B and Fig3C). Under both these conditions NPCs are clustered. However, whether any post translational modification of nucleoporins such as phosphorylation or ubiquitination facilitates such re-localisation of Uip4 is a possibility that needs to be investigated (Saleem et al., 2010; Smoyer et al., 2019).

The protein constituents of NPCs, which are among the longest lived protein complexes of the cell, undergo turnover. Regulation of starvation and aging induced nucleoporin turnover is of particular interest in the field of neurodegenerative diseases. Recent work has contributed to our understanding of how NE integrity and NPC turnover is regulated under specific conditions such as either nitrogen or carbon starvation (Tomioka et al., 2020; Lee et al., 2020). Such response is brought about by a coordinated action of players from multiple organelles. Since Uip4 bears a E-set domain that senses AMPK levels, it is likely to function in coordination with pathways involved in regulation of nutrient sensing to NE homeostasis. Why nuclear shape and NPC distribution is responsive to growth conditions is not known. However, the regulation of transcription at the nuclear periphery and association of mRNAs with the NPCs might have a role in this. Which components directly mediate the response to nutrient availability to the nuclear periphery and dictate shape alterations is an open question. Together, this study hints at the key role of a NE/ER protein in nuclear structure and function regulation under altered metabolic conditions. Future experiments will aim to identify the underlying pathways that control nuclear morphology and its components in response to such physiologically relevant signals.

## Acknowledgement

Authors would like to thank Dr Naresh Kumar M (University of Hyderabad, Hyderabad) and Dr. Sharada Sawant (Officer-In-Charge, Electron Microscopy Facility, Advanced Center for Treatment, Research & Education in Cancer, Tata Memorial Center, Mumbai) for TEM. Prof. T.C.Nag, Prof. S.C. Yadav and staff of SAIF, AIIMS, New Delhi are also acknowledged for TEM support. We would like to acknowledge Rolf Sternglanz, Karsten Weis and Valerie Doye for plasmids and C. Patrick Lusk for strains.

## Author contributions

PD contributed to the acquisition of data, analysis, and interpretation of data. IJ established the genome-wide screen, generated constructs and performed some of the imaging and analysis. KM contributed to the conception and design, analysis and interpretation of data, and acquisition of funding. KM and PD wrote the manuscript. All authors read and approved the final manuscript.

## Declaration

The authors declare that they have no competing interests.

## References

Abraham, N.M., K. Ramalingam, S. Murthy, and K. Mishra. 2019. Siz2 Prevents Ribosomal DNA Recombination by Modulating Levels of Tof2 in Saccharomyces cerevisiae. mSphere. 4. doi:10.1128/mSphere.00713-19.

Aitchison, J.D., and M.P. Rout. 2012. The Yeast Nuclear Pore Complex and Transport Through It. Genetics. 190:855–883. doi:10.1534/genetics.111.127803.

Alber, F., S. Dokudovskaya, L.M. Veenhoff, W. Zhang, J. Kipper, D. Devos, A. Suprapto, O. Karni-Schmidt, R. Williams, B.T. Chait, A. Sali, and M.P. Rout. 2007. The molecular architecture of the nuclear pore complex. Nature. 450:695–701. doi:10.1038/nature06405.

Allegretti, M., C.E. Zimmerli, V. Rantos, F. Wilfling, P. Ronchi, H.K.H. Fung, C.-W. Lee, W. Hagen, B. Turoňová, K. Karius, M. Börmel, X. Zhang, C.W. Müller, Y. Schwab, J. Mahamid, B. Pfander, J. Kosinski, and M. Beck. 2020. In-cell architecture of the nuclear pore and snapshots of its turnover. Nature. 586:796–800. doi:10.1038/s41586-020-2670-5.

Beck, M., and E. Hurt. 2016. The nuclear pore complex: understanding its function through structural insight. Nat. Rev. Mol. Cell Biol. 18:73–89. doi:10.1038/nrm.2016.147.

Bucci, M., and S.R. Wente. 1998. A novel fluorescence-based genetic strategy identifies mutants of Saccharomyces cerevisiae defective for nuclear pore complex assembly. Mol. Biol. Cell. 9:2439–61.

Casey, A.K., S. Chen, P. Novick, S. Ferro-Novick, and S.R. Wente. 2015. Nuclear pore complex integrity requires Lnp1, a regulator of cortical endoplasmic reticulum. Mol. Biol. Cell. 26. doi:10.1091/mbc.E15-01-0053.

Colombi, P., B.M. Webster, F. Fröhlich, and C.P. Lusk. 2013. The transmission of nuclear pore complexes to daughter cells requires a cytoplasmic pool of Nsp1. J. Cell Biol. 203:215–32. doi:10.1083/jcb.201305115.

Dawson, T.R., M.D. Lazarus, M.W. Hetzer, and S.R. Wente. 2009. ER membrane-bending proteins are necessary for de novo nuclear pore formation. J. Cell Biol. 184:659–75. doi:10.1083/jcb.200806174.

Deolal, P., and K. Mishra. 2021. Regulation of diverse nuclear shapes : pathways working independently, together. Commun. Integr. Biol. 14:158–175. doi:10.1080/19420889.2021.1939942.

Doye, V., R. Wepf, and E.C. Hurt. 1994. A novel nuclear pore protein Nup133p with distinct roles in poly(A)+ RNA transport and nuclear pore distribution. EMBO J. 13:6062–75.

Fernandez-Martinez, J., and M.P. Rout. 2009. Nuclear pore complex biogenesis. Curr. Opin. Cell Biol. 21:603–12. doi:10.1016/j.ceb.2009.05.001.

Flemming, D., P. Sarges, P. Stelter, A. Hellwig, B. Böttcher, and E. Hurt. 2009. Two structurally distinct domains of the nucleoporin Nup170 cooperate to tether a subset of nucleoporins to nuclear pores. J. Cell Biol. 185:387–395. doi:10.1083/jcb.200810016.

Garapati, H.S., and K. Mishra. 2018. Comparative genomics of nuclear envelope proteins. BMC Genomics. 19:823. doi:10.1186/s12864-018-5218-4.

Gardner, J.M., and S.L. Jaspersen. 2014. Manipulating the Yeast Genome: Deletion, Mutation, and Tagging by PCR. In Methods in molecular biology (Clifton, N.J.). Humana Press, New York, NY. 45–78.

Joong Kim, S., J. Fernandez-martinez, ilona Nudelman, Y. Shi, W. Zhang, B. raveh, T. herricks, B.D. Slaughter, J. hogan, P. Upla, ilan E. chemmama, riccardo Pellarin, ignacia Echeverria, manjunatha Shivaraju, azraa S. chaudhury, J. Wang, rosemary Williams, J. Unruh, charles Greenberg, E.Y. Jacobs, Z. Yu, J. de la cruz, roxana mironska, D.L. Stokes, J.D. aitchison, martin F. Jarrold, J.L. Gerton, S.J. Ludtke, christopher W. akey, B.T. chait, and andrej Sali. 2018. Integrative structure and functional anatomy of a nuclear pore complex. doi:10.1038/nature26003.

Katta, S.S., C.J. Smoyer, and S.L. Jaspersen. 2014. Destination: inner nuclear membrane. Trends Cell Biol. 24:221–229. doi:10.1016/j.tcb.2013.10.006.

Kumanski, S., B. Viart, S. Kossida, and M. Moriel-Carretero. 2021. Lipid Droplets Are a Physiological Nucleoporin Reservoir. Cells. 10:472. doi:10.3390/cells10020472.

Lee, C.-W., F. Wilfling, P. Ronchi, M. Allegretti, S. Mosalaganti, S. Jentsch, M. Beck, and B. Pfander. 2020. Selective autophagy degrades nuclear pore complexes. Nat. Cell Biol. 22:159–166. doi:10.1038/s41556-019-0459-2.

Lewis, A., R. Felberbaum, and M. Hochstrasser. 2007. A nuclear envelope protein linking nuclear pore basket assembly, SUMO protease regulation, and mRNA surveillance. J. Cell Biol. 178:813–27. doi:10.1083/jcb.200702154.

Lone, M.A., A.E. Atkinson, C.A. Hodge, S. Cottier, F. Martínez-Montañés, S. Maithel, L. Mène-Saffrané, C.N. Cole, and R. Schneiter. 2015. Yeast Integral Membrane Proteins Apq12, Brl1, and Brr6 Form a Complex Important for Regulation of Membrane Homeostasis and Nuclear Pore Complex Biogenesis. Eukaryot. Cell. 14:1217–27. doi:10.1128/EC.00101-15.

Longtine, M.S., A. Mckenzie III, D.J. Demarini, N.G. Shah, A. Wach, A. Brachat, P. Philippsen, and J.R. Pringle. 1998. Additional modules for versatile and economical PCR-based gene deletion and modification in Saccharomyces cerevisiae. Yeast. 14:953–961. doi:10.1002/(SICI)1097-0061(199807)14:10<953::AID-YEA293>3.0.CO;2-U.

Lord, C.L., and S.R. Wente. 2020. Nuclear envelope–vacuole contacts mitigate nuclear pore complex assembly stress. J. Cell Biol. 219. doi:10.1083/jcb.202001165.

Makio, T., D.L. Lapetina, and R.W. Wozniak. 2013. Inheritance of yeast nuclear pore complexes requires the Nsp1p subcomplex. J. Cell Biol. 203:187–96. doi:10.1083/jcb.201304047.

Makio, T., L.H. Stanton, C.-C. Lin, D.S. Goldfarb, K. Weis, and R.W. Wozniak. 2009. The nucleoporins Nup170p and Nup157p are essential for nuclear pore complex assembly. J. Cell Biol. 185:459–73. doi:10.1083/jcb.200810029.

Male, G., P. Deolal, N.K. Manda, S. Yagnik, A. Mazumder, and K. Mishra. 2020. Nucleolar size regulates nuclear envelope shape in Saccharomyces cerevisiae. J. Cell Sci. doi:10.1242/jcs.242172.

Marchler-Bauer, A., S. Lu, J.B. Anderson, F. Chitsaz, M.K. Derbyshire, C. DeWeese-Scott, J.H. Fong, L.Y. Geer, R.C. Geer, N.R. Gonzales, M. Gwadz, D.I. Hurwitz, J.D. Jackson, Z. Ke, C.J. Lanczycki, F. Lu, G.H. Marchler, M. Mullokandov, M. V. Omelchenko, C.L. Robertson, J.S. Song, N. Thanki, R.A. Yamashita, D. Zhang, N. Zhang, C. Zheng, and S.H. Bryant. 2011. CDD: a Conserved Domain Database for the functional annotation of proteins. Nucleic Acids Res. 39:D225–D229. doi:10.1093/nar/gkq1189.

Mekhail, K., and D. Moazed. 2010. The nuclear envelope in genome organization, expression and stability. Nat. Rev. cell Biol. 11:317–328. doi:10.1038/nrm2894.

Mészáros, N., J. Cibulka, M.J. Mendiburo, A. Romanauska, M. Schneider, and A. Köhler. 2015. Nuclear Pore Basket Proteins Are Tethered to the Nuclear Envelope and Can Regulate Membrane Curvature. Dev. Cell. doi:10.1016/j.devcel.2015.02.017.

Onischenko, E., L.H. Stanton, A.S. Madrid, T. Kieselbach, and K. Weis. 2009. Role of the Ndc1 interaction network in yeast nuclear pore complex assembly and maintenance. J. Cell Biol. 185:475–91. doi:10.1083/jcb.200810030.

Otsuka, S., A.M. Steyer, M. Schorb, J.-K. Hériché, M.J. Hossain, S. Sethi, M. Kueblbeck, Y. Schwab, M. Beck, and J. Ellenberg. 2018. Postmitotic nuclear pore assembly proceeds by radial dilation of small membrane openings. Nat. Struct. Mol. Biol. 25:21–28. doi:10.1038/s41594-017-0001-9.

Rajoo, S., P. Vallotton, E. Onischenko, and K. Weis. 2018. Stoichiometry and compositional plasticity of the yeast nuclear pore complex revealed by quantitative fluorescence microscopy. Proc. Natl. Acad. Sci. U. S. A. 201719398. doi:10.1073/pnas.1719398115.

Rout, M.P., J.D. Aitchison, A. Suprapto, K. Hjertaas, Y. Zhao, and B.T. Chait. 2000. The yeast nuclear pore complex: composition, architecture, and transport mechanism. J. Cell Biol. 148:635–51. doi:10.1083/jcb.148.4.635.

Rout, M.P., and S.R. Wente. 1994. Pores for thought: nuclear pore complex proteins. Trends Cell Biol. 4:357–365. doi:10.1016/0962-8924(94)90085-X.

Rouvière, J.O., M. Bulfoni, A. Tuck, B. Cosson, F. Devaux, and B. Palancade. 2018. A SUMO-dependent feedback loop senses and controls the biogenesis of nuclear pore subunits. Nat. Commun. 9:1665. doi:10.1038/s41467-018-03673-3.

Sakuma, S., and M.A. D’Angelo. 2017. The roles of the nuclear pore complex in cellular dysfunction, aging and disease. Semin. Cell Dev. Biol. doi:10.1016/j.semcdb.2017.05.006.

Saleem, R.A., R.S. Rogers, A. V. Ratushny, D.J. Dilworth, P.T. Shannon, D. Shteynberg, Y. Wan, R.L. Moritz, A.I. Nesvizhskii, R.A. Rachubinski, and J.D. Aitchison. 2010. Integrated Phosphoproteomics Analysis of a Signaling Network Governing Nutrient Response and Peroxisome Induction. Mol. Cell. Proteomics. 9:2076–2088. doi:10.1074/mcp.M000116-MCP201.

Scarcelli, J.J., C.A. Hodge, and C.N. Cole. 2007. The yeast integral membrane protein Apq12 potentially links membrane dynamics to assembly of nuclear pore complexes. J. Cell Biol. 178.

Schneiter, R., and C.N. Cole. 2010. Integrating complex functions: coordination of nuclear pore complex assembly and membrane expansion of the nuclear envelope requires a family of integral membrane proteins. Nucleus. 1:387–92. doi:10.4161/nucl.1.5.12333.

Smoyer, C.J., S.E. Smith, J.M. Gardner, S. McCroskey, J.R. Unruh, and S.L. Jaspersen. 2019. Distribution of Proteins at the Inner Nuclear Membrane Is Regulated by the Asi1 E3 Ligase in Saccharomyces cerevisiae. Genetics. 211:1269–1282. doi:10.1534/genetics.119.301911.

Stade, K., C.S. Ford, C. Guthrie, and K. Weis. 1997. Exportin 1 (Crm1p) is an essential nuclear export factor. Cell. 90:1041–50.

Strunov, A.A., E.A. Onishchenko, and E. V. Kiseleva. 2011. Inhibition of POM152 and POM34 expression in budding yeast arrests the assembly of the nuclear pore complex and increases its inner diameter. Russ. J. Genet. Appl. Res. doi:10.1134/S2079059711030117.

Takahashi, Y., J. Mizoi, and A. Tohe. 2000a. Yeast Ulp1, an Smt3-Specific Protease, Associates with Nucleoporins1. Rapid Commun. J. Bi ochem. 128:723–725.

Takahashi, Y., J. Mizoi, A. Toh-e, and Y. Kikuchi. 2000b. Yeast Ulpl, an Smt3-Specific Protease, Associates with Nucleoporins. J. Biochem. 128:723–725. doi:10.1093/oxfordjournals.jbchem.a022807.

Titus, L.C., T.R. Dawson, D.J. Rexer, K.J. Ryan, and S.R. Wente. 2010. Members of the RSC chromatin-remodeling complex are required for maintaining proper nuclear envelope structure and pore complex

Tirupataiah, S., I. Jamir, I. Srividya, and K. Mishra. 2014. Yeast Nkp2 is required for accurate chromosome segregation and interacts with several components of the central kinetochore. Mol. Biol. Rep. 41. doi:10.1007/s11033-013-2918-3.

Tomioka, Y., T. Kotani, H. Kirisako, Y. Oikawa, Y. Kimura, H. Hirano, Y. Ohsumi, and H. Nakatogawa. 2020. TORC1 inactivation stimulates autophagy of nucleoporin and nuclear pore complexes. J. Cell Biol. 219. doi:10.1083/JCB.201910063.

Webster, M., K.L. Wikin, and O. Cohen-Fix. 2009. Sizing up the nucleus: Nuclear shape, size and nuclear-envelope assembly. J. Cell Sci. doi:10.1242/jcs.037333

Webster, B.M., P. Colombi, J. Jäger, and C.P. Lusk. 2014. Surveillance of nuclear pore complex assembly by ESCRT-III/Vps4. Cell. 159:388–401. doi:10.1016/j.cell.2014.09.012.

Wente, S.R., and G. Blobel. 1993. A temperature-sensitive NUP116 null mutant forms a nuclear envelope seal over the yeast nuclear pore complex thereby blocking nucleocytoplasmic traffic. J. Cell Biol. 123:275–284. doi:10.1083/jcb.123.2.275.

Yewdell, W.T., P. Colombi, T. Makhnevych, and C.P. Lusk. 2011. Lumenal interactions in nuclear pore complex assembly and stability. Mol. Biol. Cell. 22:1375–88. doi:10.1091/mbc.E10-06-0554.

