## Supplemental figures for "Uip4p modulates nuclear pore complex function in *Saccharomyces cerevisiae*"

**Supplementary figures:**

### Figure S1

A

| Systematic Name | Yeast Gene Name | Mammalian Counterpart | Nuclear envelope defect | Other known function in yeast |
|---|---|---|---|---|
| YDR192C | UIP1, NUP42, RIP1 | Human NUP42 can complement yeast mutant | Mild | NPC related functions |
| YAL014C | UIP2, SYN8, SLT2 | STX8- Syntaxin 8 | Moderate | Involved in Golgi to Vacuole transport |
| YAR027W | UIP3 | None | Severe | Unknown |
| YPL186C | UIP4 | None | Severe | Unknown |
| YKR044W | UIP5 | LMAN2- Lectin mannose binding 2 | Mild | Predicted to function in Golgi and ER organization |

C

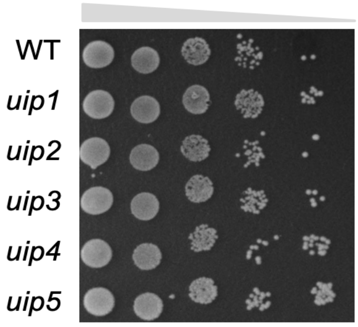


B

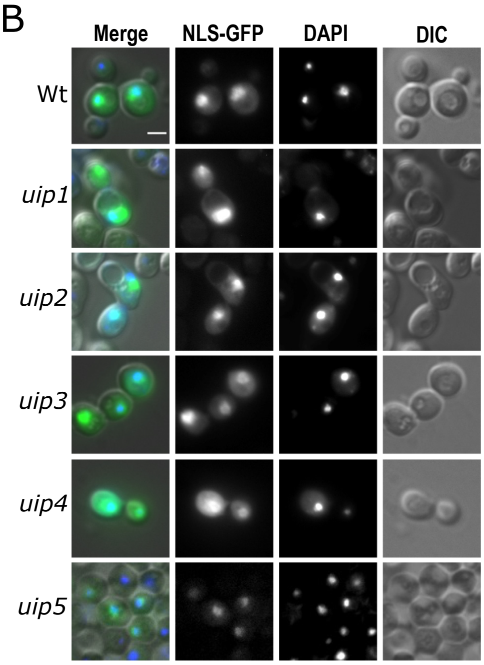


D

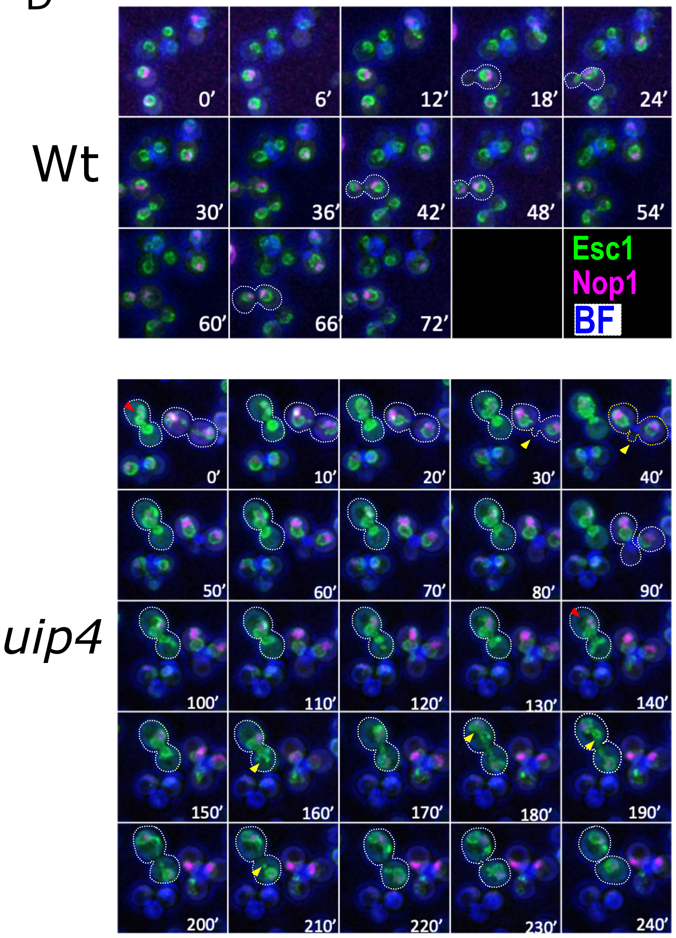


E

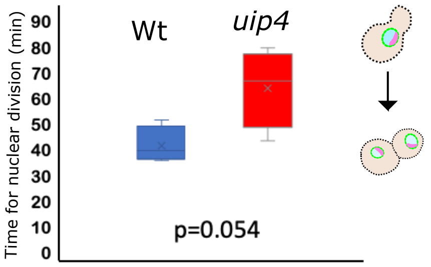


F

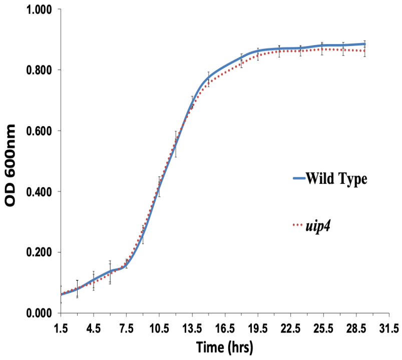

### Figure S1

FigureS1.
A. Table with detail of UIPs screened for nuclear shape defect is presented. Information presented is sourced from Saccharomyces Genome Database.
B. Nuclear import was tested in Wt and indicated strains bearing NLS-2X GFP plasmid. DAPI staining was used to define the nucleus. Scale-2μm
C. In order to assess growth phenotype, overnight cultures of the indicated strains were taken and sub-cultured by inoculating an equal number of cells in a fresh medium for 4 hours. Cells were then harvested and 10-fold dilutions were serially spotted on a SC-plate. The plates were incubated at 30ºC for 2 days prior to imaging.
D. MIP of Wt and uip4 marking NE (GFP-Esc1) in green and nucleolus (mRFP-Nop1) in magenta acquired at indicated time points during time lapse live cell imaging are shown. Arrow heads- associated NE abnormality.
E. The plot indicates the time taken by Wt and uip4 cells to complete nuclear division. Horizontal line represents the mean.
F. Overnight cultures of the indicated strains were taken and sub-cultured into fresh medium at 30ºC. OD600 was recorded every 90 min and plotted. Three biological replicates of Wt (blue) and uip4 (red) cells were used. Error bars represent SEM.

### Figure S2

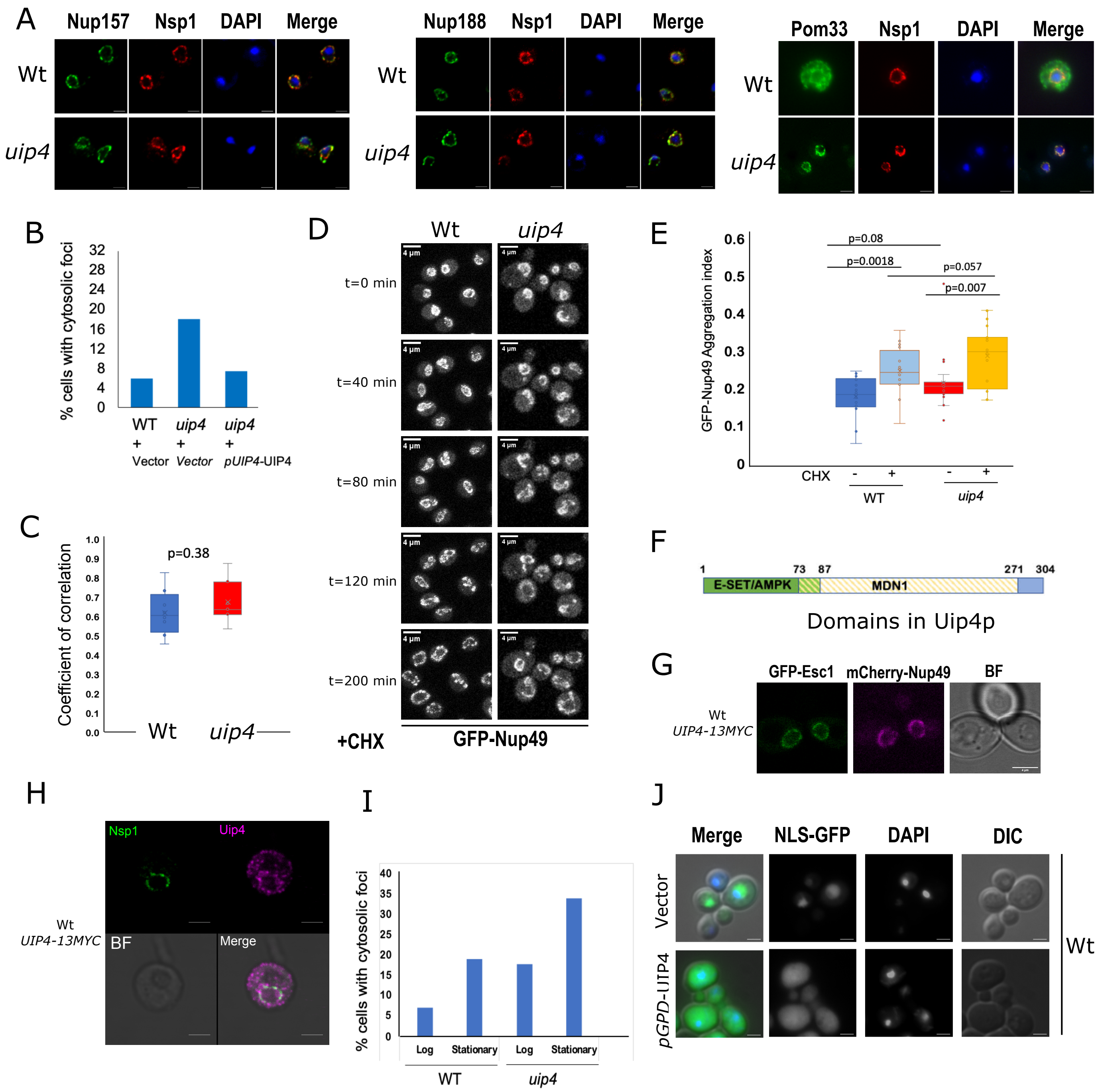


**FigureS2.**

A. Indirect immunofluorescence was performed in strains expressing either Nup157, Nup188 or Pom33 tagged with GFP. GFP and Nsp1 antibodies were used to detect respective nucleoporins. DAPI is used as a nuclear stain. Scale-2µm.

B. The bar graph represents the fraction of cells in the indicated strain complemented with either empty vector or vector with UIP4 cloned with its native promoter, showing cytosolic spots of GFP-Nup49. ~100 cells from 2 independent experiments were counted.

C. The extent of co-localization for Nup157-GFP and Nup49- mCherry shown in Fig2D is assessed by measuring the correlation coefficient between the two signals. Horizontal line represents the mean.

D. MIP of Wt and uip4 expressing GFP-Nup49 acquired during time lapse live cell imaging are shown. Time indicates minutes after cycloheximide (CHX) addition. Scale-4µm

E. The box plot represents the aggregation index calculated from GFP signal along the NE in the mid-focal plane of 20-25 individual cells as indicated. Horizontal line is the mean value.

F. Domain map of Uip4 showing position of E-set AMPK and the MDN1 domain.

G. Strain expressing Uip4 tagged with 13xMyc epitope at endogenous loci was co-transformed with plasmids expressing GFP-Esc1 and Nup49-mCherry. Live cells were imaged and representative images are shown. Scale-2µm

H. Indirect immunofluorescence using α-myc was performed in the strain carrying UIP4-13MYC to check the localization of Uip4. Co-staining with a NE marker Nsp1 is shown. Scale 2µm

I. The bar graph represents the fraction of cells in the indicated strain harvested from either mid-log or stationary phase showing cytosolic spots of GFP-Nup49. ~100 cells from 2 independent experiments were counted.

J. Nuclear import was tested in Wt cells bearing NLS-2X GFP plasmid, co-transformed with either empty vector or vector expressing UIP4 from pGPD. DAPI staining was used to define the nucleus. Scale-2µm
